# TeoNAM: a nested association mapping population for domestication and agronomic trait analysis in maize

**DOI:** 10.1101/647461

**Authors:** Qiuyue Chen, Chin Jian Yang, Alessandra M. York, Wei Xue, Lora L. Daskalska, Craig A. DeValk, Kyle W. Krueger, Samuel B. Lawton, Bailey G. Spiegelberg, Jack M. Schnell, Michael A. Neumeyer, Joseph S. Perry, Aria C. Peterson, Brandon Kim, Laura Bergstrom, Liyan Yang, Isaac C. Barber, Feng Tian, John F. Doebley

## Abstract

Recombinant inbred lines (RILs) are an important resource for mapping genes controlling complex traits in many species. While RIL populations have been developed for maize, a maize RIL population with multiple teosinte inbred lines as parents has been lacking. Here, we report a teosinte nested association mapping population (TeoNAM), derived from crossing five teosinte inbreds to the maize inbred line W22. The resulting 1257 BC_1_S_4_ RILs were genotyped with 51,544 SNPs, providing a high-density genetic map with a length of 1540 cM. On average, each RIL is 15% homozygous teosinte and 8% heterozygous. We performed joint linkage mapping (JLM) and genome-wide association study (GWAS) for 22 domestication and agronomic traits. A total of 255 QTLs from JLM were identified with many of these mapping to known genes or novel candidate genes. TeoNAM is a useful resource for QTL mapping for the discovery of novel allelic variation from teosinte. TeoNAM provides the first report that *PROSTRATE GROWTH1*, a rice domestication gene, is also a QTL associated with tillering in teosinte and maize. We detected multiple QTLs for flowering time and other traits for which the teosinte allele contributes to a more maize-like phenotype. Such QTL could be valuable in maize improvement.

## Introduction

Recombinant inbred line (RIL) populations are powerful tools for investigating the genetic architecture of traits and identifying the causal genes that underlie trait variation. RIL populations have been widely used in many organisms. In mammals, the well-known Collaborative Cross (CC), consisting of a large panel of multiparental recombinant inbred mouse lines, has been specifically designed for the analysis of complex traits (Churchill *et al*. 2004). Similarly, the Drosophila Synthetic Population Resource (DSPR), which consists of two sets of RILs, has been designed to combine the high mapping resolution offered by multiple generations of recombination with the high statistical power afforded by a linkage-based design (King *et al*. 2012). In plants, the maize nested association mapping population (NAM), which crossed 25 founders to a common parent in maize (Yu *et al*. 2008), has been successfully applied to a large number of traits (Buckler *et al*. 2009; Tian *et al*. 2011; Kump *et al*. 2011). The NAM design has also been utilized to other crops such as barley (Maurer *et al*. 2015; Nice *et al*. 2016), rice (Fragoso *et al*. 2017), sorghum (Bouchet *et al*. 2017), wheat (Jordan *et al*. 2018), and soybean (Xavier *et al*. 2018). In Arabidopsis, another design, called Multiparent Advanced Generation Inter-Cross (MAGIC) population, provides high precision to detect QTLs (Kover *et al*. 2009; Huang *et al*. 2011). This design has also been used in wheat (Huang *et al*. 2012; Mackay *et al*. 2014), rice (Bandillo *et al*. 2013), and maize (Dell’Acqua *et al*. 2015; Xiao *et al*. 2016).

For the study of maize domestication, many new discoveries were made using a biparental maize-teosinte BC_2_S_3_ RIL population. Shannon (2012) performed QTL mapping for 16 traits and examined the genetic architecture of domestication at the whole genome level. This RIL population has also been widely used to fine-map QTL and identify causal or candidate genes for many QTLs including ones controlling seed shattering (Lin *et al*. 2012), leaf number (Li *et al*. 2016), kernel row number (Calderón *et al*. 2016), shoot apical meristem morphology (Leiboff *et al*. 2016), vascular bundle number (Huang *et al*. 2016), tassel related traits (Xu *et al*. 2017b), and nodal root number (Zhang *et al*. 2018). With this population, several QTL have been fine-mapped to single genes including *grassy tillers1* (*gt1*) for controlling prolificacy (Wills *et al*. 2013), *prolamin-box binding factor1* (*pbf1*) for kernel weight (Lang *et al*. 2014), *glossy15* (*gl15*) for vegetative phase changes (Xu *et al*. 2017a), *ZmCCT10* for photo-period response (Hung *et al*. 2012), and *zea agamous-like1* (*zagl1*) for kernel row number and flowering time (Wills *et al*. 2017), as well as several more genes regulating flowering time: *ZmCCT9* (Huang *et al*. 2018), *Zea mays CENTRORADIALIS8* (*ZCN8)* (Guo *et al*. 2018), and *ZmMADS69* (Liang *et al*. 2018). In addition to phenotypic traits, the maize-teosinte BC_2_S_3_ RIL population was used for a comprehensive genome-wide eQTL analysis to study the changes in gene expression during maize domestication (Wang *et al*. 2018).

Despite its utility, the maize-teosinte BC_2_S_3_ RIL population has three limitations. First, there is only a single teosinte parent, which cannot broadly represent the diversity of teosinte. Second, this population had two generations of backcross, which produces a background in which some teosinte traits are suppressed and do not segregate among the RILs. Third, the teosinte parent was a wild outcrossed individual which, unlike an inbred line, could not be maintained as a permanent resource.

In this paper, we report the development of a teosinte NAM population (TeoNAM) of 1257 BC_1_S_4_ RILs using five teosinte inbred parents crossed with a common maize parent (W22) for mapping QTLs for domestication and agronomic traits. We have genotyped the RILs with 51,544 genotype-by-sequencing (GBS) markers that provide a high-density genetic map. The TeoNAM population captures a large number of recombination events for localizing QTL to genomic locations and the single generation of backcross allows enhanced expression of teosinte traits as compared to the BC_2_S_3_ RIL population. We report data for 22 traits but focus our discussion on 9 traits to illustrate the utility of TeoNAM including identifying candidate genes. TeoNAM will be a valuable resource for dissecting the genetic basis of domestication and agronomic traits.

## Results

### Characterization of a teosinte NAM population

We developed a teosinte NAM population (TeoNAM), which was constructed by crossing five teosinte inbred lines to a maize inbred line W22, followed by one generation of backcross to the common recurrent maize parent and four generations of selfing (Figure S1). The teosinte parents include four *Zea mays* ssp. *parviglumis* lines and one *Zea mays* ssp. *mexicana* line. As such, TeoNAM encompasses five bi-parental families, each with 219-310 BC_1_S_4_-derived recombinant inbred lines (RILs) for a total of 1257 RILs. The number of segregating SNP markers range from 11,395 to 16,109 per family with over 51,000 total SNP markers (Table 1).

**Table 1.** TeoNAM genetic map statistics.

| **Table 1. TeoNAM genetic map statistics** |  |  |  |  |  |  |  |  |
| --- | --- | --- | --- | --- | --- | --- | --- | --- |
| **Population** | **No.**<br>**RILs** | **No.**<br>**Markers** | **Length**<br>**(cM)** | **No.**<br>**XOs** | **cM/Mb** | **W22**<br>**(%)** | **Heterozygous**<br>**(%)** | **Teosinte**<br>**(%)** |
| W22×TIL01 | 223 | 13,088 | 1457 | 6,291 | 0.71 | 75.8 | 7.7 | 16.0 |
| W22×TIL03 | 270 | 16,109 | 1596 | 8,505 | 0.78 | 75.5 | 8.1 | 16.2 |
| W22×TIL11 | 219 | 13,187 | 1398 | 5,745 | 0.68 | 76.3 | 7.6 | 15.6 |
| W22×TIL14 | 235 | 11,395 | 1348 | 6,462 | 0.65 | 75.7 | 9.4 | 14.6 |
| W22×TIL25 | 310 | 14,884 | 1506 | 8,877 | 0.73 | 77.6 | 8.0 | 14.2 |
| Composite | 1257 | 51,544 | 1540 | 35,880 | 0.75 | 76.6 | 8.1 | 15.0 |

The expected segregation for a BC_1_S_4_ population is 73.44% homozygous recurrent, 3.13% heterozygous, and 23.44% homozygous donor parent. Overall, the percentage of genotypes observed were 76.6% W22 homozygous, 15% teosinte homozygous and 8.1% heterozygous across all SNP sites in the TeoNAM population (Table 1). The percentage of teosinte varied among subpopulations from 14.2%-16.2% (Table 1) and also varied across the genome in all subpopulations (Figure S2). The observed higher than expected heterozygosity may be due to unconscious selection for more heterozygous plants which had hybrid vigor. The chromosomal region of highest heterozygosity is on the short arm of chromosome 4 near *teosinte glume architecture1* (*tga1*) (Wang *et al.* 2005). Selection against homozygotes for the teosinte allele of *tga1*, which have poor ear and kernel quality, may be the cause. For a BC_1_S_4,_ the expected frequency of the maize allele is 75%. All subpopulations deviate from this with an excess of maize allele (Table 1) and the amount of excess varies across the genome (Figure S3).

We constructed genetic linkage maps for each family and a composite linkage map based on all RILs across all families and identified and annotated 51,544 high confidence SNPs that were used to impute the SNP alleles in the RILs. The composite genetic map based on these markers is 1540 cM in length including 35,880 crossovers. We examined the relationship between genetic distance in cM and physical distance in Mb based on the composite genetic map. The mean value is 0.75 cM/Mb. However, there is a wide deviation from the mean across the genome (0 - 5.52 cM/Mb). As expected, there is suppressed recombination near the centromeres (Figure S2) and frequent recombination near the telomeres where gene density is high as well (Figure S2).

We scored 22 traits for the TeoNAM lines of which 15 traits are domestication related, including vegetative gigantism (CULM, LFLN, and LFWD), prolificacy (PROL), tillering (TILN), ear shattering (SHN), conversion of the inflorescence from staminate to pistillate (STAM), multiple ear-related traits (EB, ED, EL, KRN, KW), glume traits (GLCO and GLUM), and red pericarp color (REPE) (Table 2). Additionally, several agronomic traits were scored including flowering (ASI, DTA, and DTS), plant architecture (PLHT and TBN), barren ear base (BARE), and yellow pericarp color (YEPE). Most traits (ASI, CULM, DTA, DTS, ED, EL, KRN, KW, LFLN, LFWD, PHLT, and TBN) follow approximately normal distribution, suggesting an oligo- or polygenic genetic control of these traits, but other traits (BARE, EB, GLCO, GLUM, PROL, REPE, SHN, STAM, TILN, and YEPE) exhibited a skewed or non-normal distribution. Some of these traits are meristic or discrete traits (e.g. PROL or TILN). A few traits, like STAM, show a two-part distribution with a spike at 0 plus continuous range of values from 0 to 2, which suggest they may be polygenic threshold traits (Figure S4). There are also substantial differences in trait mean among the five subpopulations, indicating underlying differences in genetic architecture among the five teosinte inbreds (Figure S5).

**Table 2.** List of 22 domestication and agronomic traits scored.

| **Table 2. List of 22 domestication and agronomic traits scored** |  |  |  |
| --- | --- | --- | --- |
| **Trait** | **Abbreviation** | **Units** | **Category** |
| Anthesis-Silk Interval | ASI | count | Agronomic |
| Barren Ear Base | BARE | score | Agronomic |
| Days to Anthesis | DTA | count | Agronomic |
| Days to Silk | DTS | count | Agronomic |
| Plant Height | PLHT | cm | Agronomic |
| Tassel Branch Number | TBN | count | Agronomic |
| Yellow Pericarp | YEPE | score | Agronomic |
| Culm Diameter | CULM | mm | Domestication |
| Ear Branch Number | EB | count | Domestication |
| Ear Diameter | ED | mm | Domestication |
| Ear Length | EL | cm | Domestication |
| Glume Color | GLCO | score | Domestication |
| Glume Score | GLUM | score | Domestication |
| Kernel Row Number | KRN | count | Domestication |
| Kernel Weight | KW | g | Domestication |
| Leaf Length | LFLN | cm | Domestication |
| Leaf Width | LFWD | cm | Domestication |
| Prolificacy | PROL | binary | Domestication |
| Red Pericarp | REPE | score | Domestication |
| Shattering | SHN | count | Domestication |
| Staminate Spikelet | STAM | score | Domestication |
| Tiller Number | TILN | count | Domestication |

### QTL mapping

We used both Joint Linkage Mapping (JLM) and the Genome-Wide Association Study (GWAS) method as two complementary approaches for QTL detection. We also used basic interval QTL mapping for the five individual subpopulations to provide a guide for future work to fine-map the genes underlying the QTL. We detected 255 QTLs for 22 traits by JLM which combines information across all families (Figure 1; Table S1). We detected a total of 150 QTLs by GWAS, among which 57 QTLs overlapped with QTLs by JLM (Table S2). Separate QTL mapping for each subpopulation detected 464 QTLs in total, among which 310 QTLs overlapped with QTLs by JLM (Figure S6-S27; Table S3). Below, we focused on QTL detected by JLM for our characterization of the genetic architecture and the distribution of QTL allelic effects.

**Figure 1.**
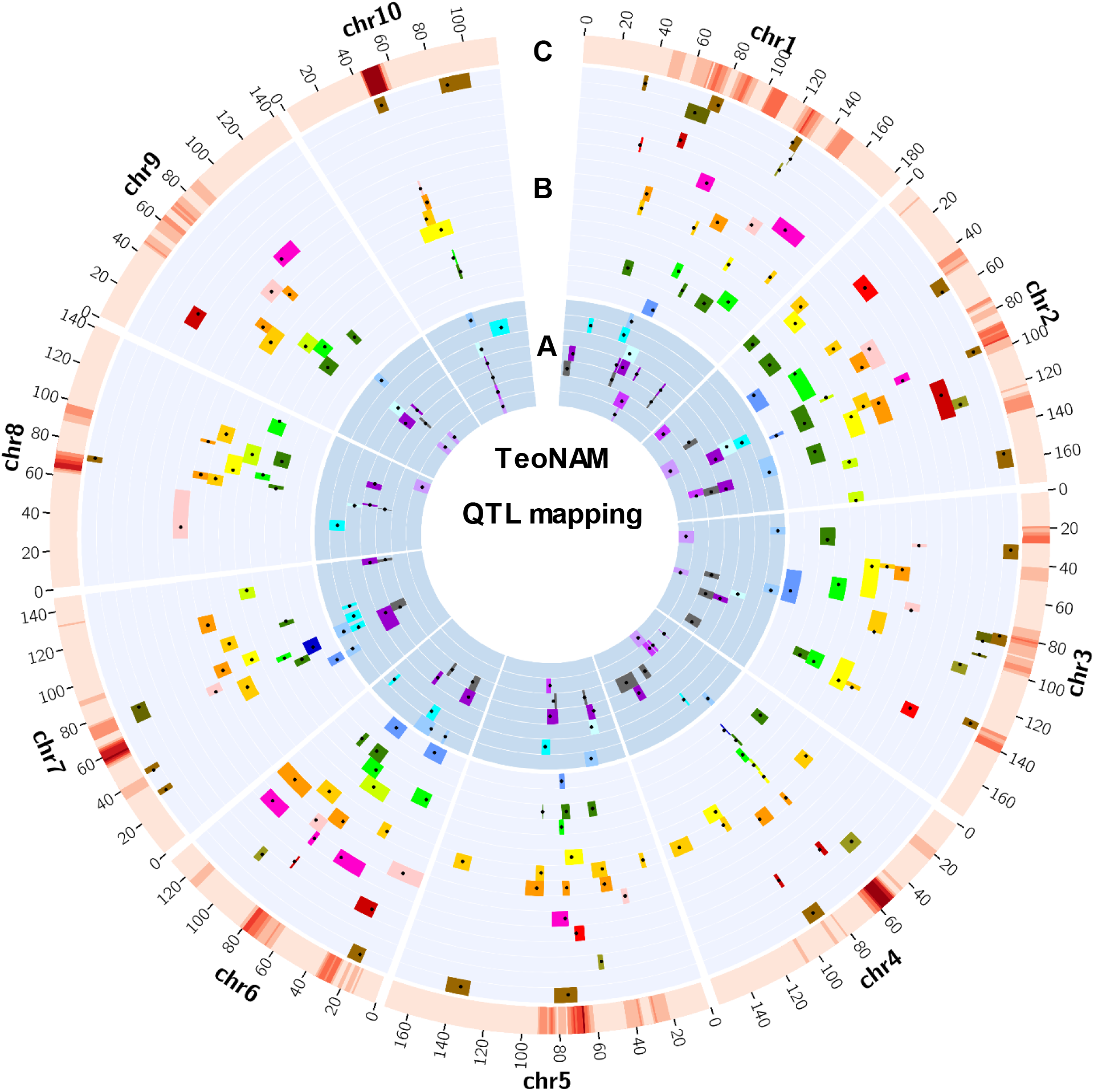
Genomic distribution of QTLs for all 22 traits in TeoNAM. The 22 agronomic and domestication (B) traits are plotted in layers with different background colors, following the order of ASI, BARE, DTA, DTS, PLHT, TBN, YEPE, CULM, EB, ED, EL, GLCO, GLUM, KRN, KW, LFLN, LFWD, PROL, REPE, SHN, STAM and TILN outwards. Black dots indicate QTL peaks detected by JLM and colored bars indicate the support interval of QTLs for different traits. The heat map in the outmost layer (C) shows the number of QTL peaks using a sliding window of 10 cM and 1 cM steps, where low to high density of QTLs (0-12) are shown in light to dark red, respectively.

Among 22 traits, the number of QTL ranges from 2 to 24; the trait with most QTL is KRN. Genetic architecture varies considerably among traits (Figure 2; Figure S28). Several traits, including BARE, GLCO, GLUM, PROL, REPE, STAM and YEPE, had relatively simple genetic architectures with two to ten QTL including one of large effect. The largest QTL for each of these traits has between 2.1 and 11.7 times the additive effect of the second largest QTL. A second class of traits have a genetic architecture that is either more polygenic (ED, KRN, KW, LFLN, TBN, and TILN) or having only a few QTL of small effect (ASI, CULM, EB, LFWD, PLHT and SHN). For these traits, there was no single large effect QTL that accounts for the majority of the explainable variation. The largest effect QTL for each of these traits has between 1 and 1.8 times the size of the effect of the second largest QTL. A final class of traits has a genetic architecture with both a single QTL of large effect plus multiple QTLs of small effect. These traits include DTA, DTS, and EL. The largest effect QTL for each of these traits is between 2.4 and 3.7 times the size of the second largest QTL.

**Figure 2.**
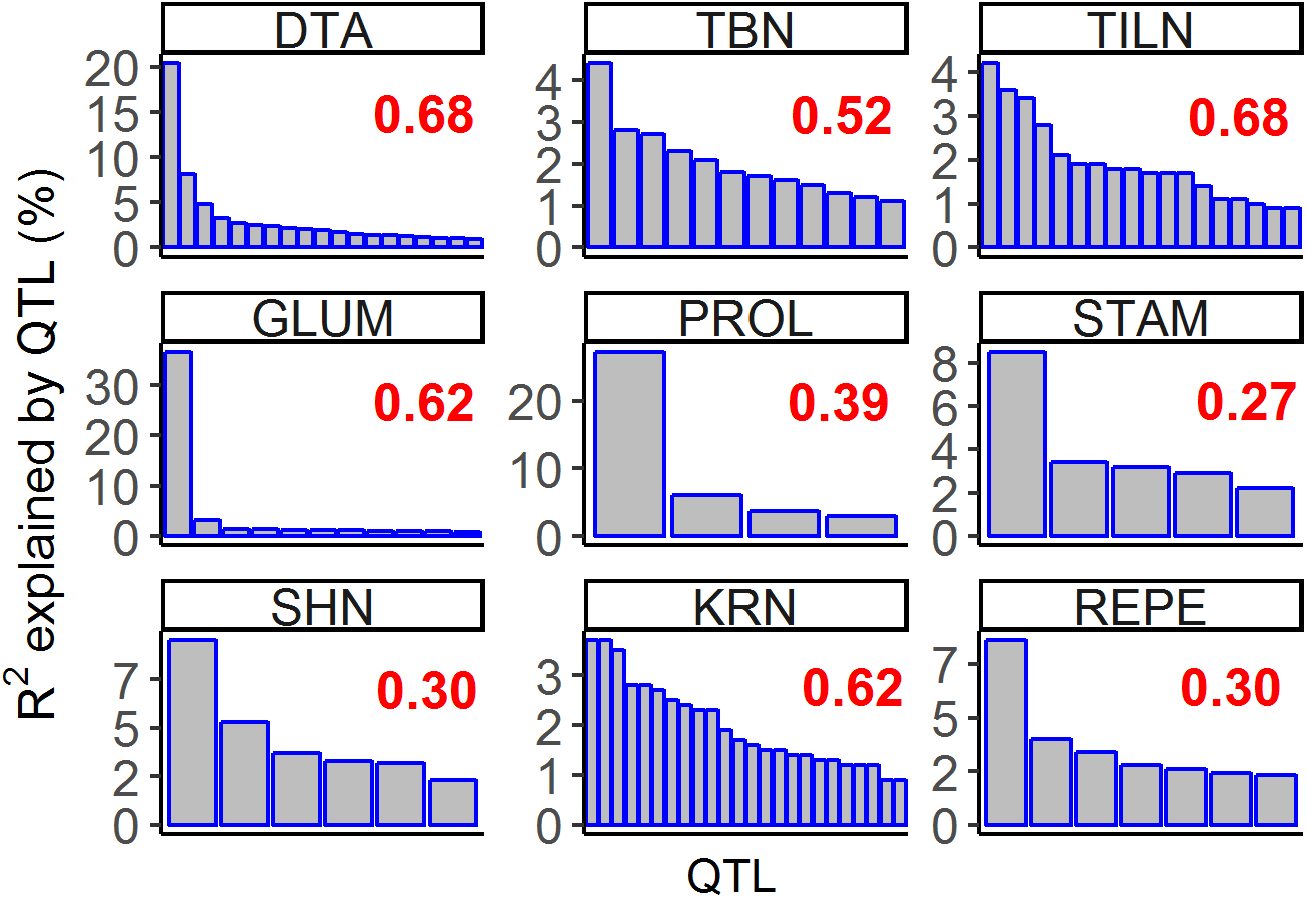
Distinct genetic architectures for different traits. The nine traits that we focused in the main text are shown. The horizontal axis indicates QTLs and the vertical axis indicates the phenotypic variation explained by each QTL (R^2^). Red number indicates variance explained by the QTL model for each trait. The R2 distribution for 13 additional traits can be found in Figure S28.

#### QTL for agronomic traits

DTA is a classical quantitative trait for maize, and in TeoNAM, it is controlled by a large-effect QTL plus many small-effect QTLs from JLM results. We detected 19 QTLs that explained 68% of the total variance for DTA (Figure 3). Among them, several recently cloned flowering time genes were detected. For example, QTL *DTA1.1* mapped to *zea agamous like1* (*zagl1*), which affects flowering time as well as multiple traits related to ear size with the maize allele conferring larger ears with more kernels (Wills *et al*. 2017). The QTL *DTA3.1* mapped to *MADS-box transcription factor69* (*ZmMADS69*), which functions as a flowering activator through the *ZmRap2.7*-*ZCN8* regulatory module and contributes to both long-day and short-day adaptation (Liang *et al*. 2018). QTL *DTA8.1* mapped to *ZCN8*, which is the maize florigen gene and has a central role in mediating flowering (Meng *et al*. 2011; Guo *et al*. 2018). QTL *DTA9.1* mapped to *ZmCCT9*, in which a distant Harbinger-like transposon acts as a *cis*-regulatory element to repress its expression to promote flowering under the long days of higher latitudes (Huang *et al*. 2018). QTL *DTA10.1* mapped to *ZmCCT10*, a known gene involved in photoperiod response in maize (Hung *et al*. 2012; Yang *et al*. 2013).

**Figure 3.**
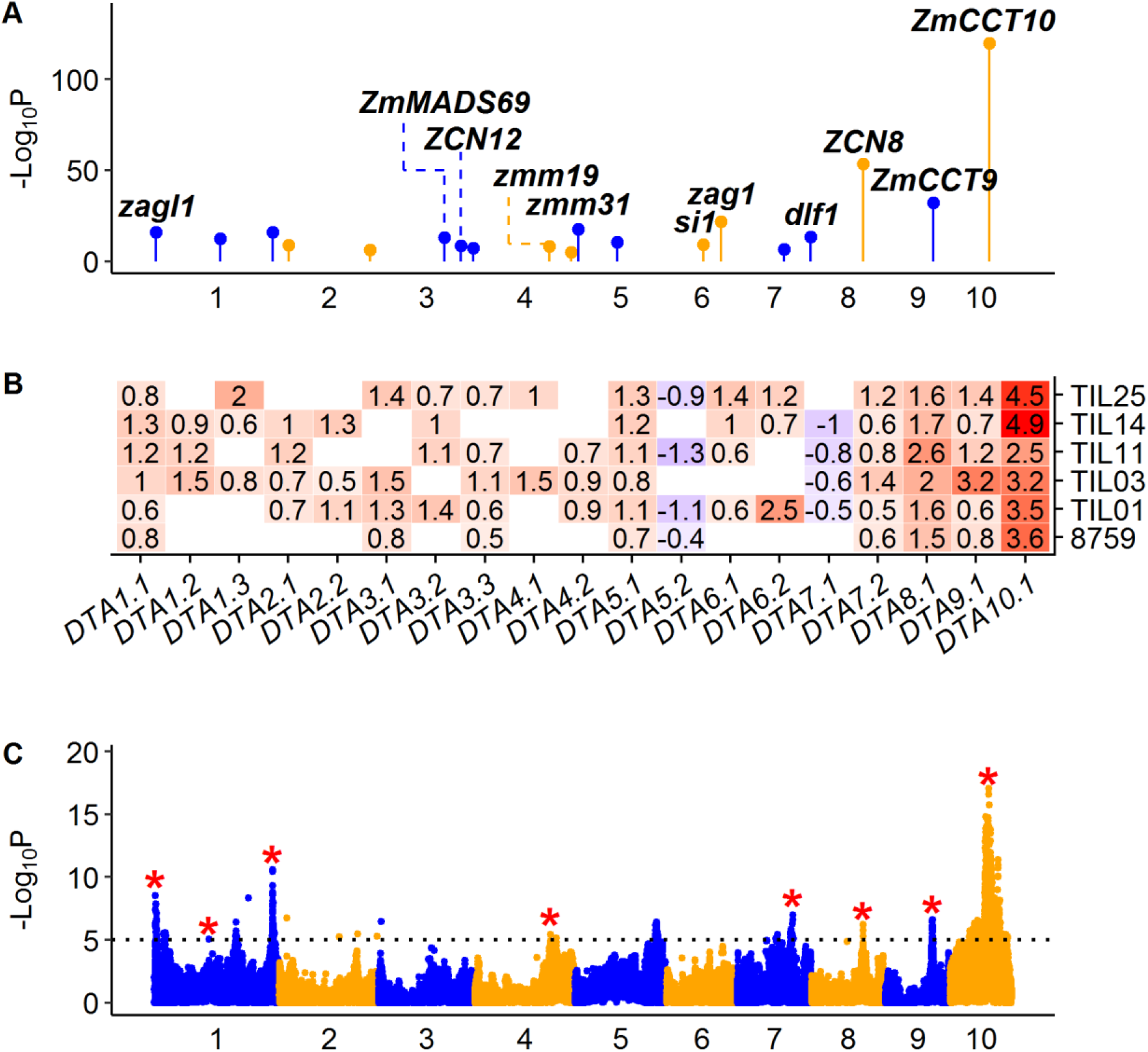
QTL characterization for agronomic trait DTA. (A) Genomic distribution of 19 QTLs for DTA detected by JLM. The known candidate genes are shown above the corresponding QTLs in bold italic. (B) Heat map shows additive allele effects of teosinte relative to maize in number of days for 19 QTLs detected by JLM. The allele effect of teosinte parent 8759 was estimated from the 866 maize-teosinte BC_2_S_3_ RILs (Shannon 2012). Insignificant effects are shown as blank. Red and blue color indicates that the teosinte allele delays or promotes flowering time, respectively. (C) Manhattan plot shows QTLs detected by GWAS. The significance threshold at LOD=5 is indicated by black dotted line. The red stars indicate GWAS signals overlapping with QTLs by JLM. In (A) and (C), chromosomes in odd and even numbers are shown in blue and orange colors, respectively.

In addition to these genes, we also identified several other candidate genes for DTA that have not previously been characterized as genes underlying a QTL. QTL *DTA3.2* mapped to *Zea mays CENTRORADIALIS12* (*ZCN12*), which is a potential floral activator (Meng *et al*. 2011). QTL *DTA4.1* mapped to *Zea mays MADS19* (*zmm19*) and *DTA5.1* mapped to *Zea mays MADS31* (*zmm31*). QTL *DTA6.1* mapped to *silky1* (*si1*), which is also a MADS box gene required for lodicule and stamen identity (Ambrose *et al*. 2000). QTL *DTA6.2* mapped to *zea agamous1* (*zag1*), which is known to affect maize flower development (Schmidt *et al*. 1993). It’s well known that MADS-box genes encode transcription factors that are key regulators of plant inflorescence and flower development (Theissen *et al*. 2000). Other than MADS genes, QTL *DTA7.2* mapped to *delayed flowering1* (*dlf1*), a floral activator gene downstream of *ZCN8* (Meng *et al*. 2011).

As expected, the teosinte alleles delayed flowering for the QTL that mapped to candidate genes. We plotted the phenotypic difference in DTA between teosinte and maize across the whole genome, and the teosinte genotype is associated with late flowering over most of the genome, even where no QTL were detected, suggesting that there are many additional minor-effect QTLs that were not detected due to insufficient statistical power (Figure S29). Interestingly, chromosome 5 and 7 are exceptions to this pattern with teosinte genotype being associated with early flowering at most sites (Figure S29). Results for DTS are similar to DTA as expected (Figure S30).

TBN is the only tassel trait we scored. We detected 12 QTLs of small effects that explained 52% of the total variance for TBN (Figure S31). Among them, several classical genes were identified. QTL *TBN6.1* mapped to *fasciated ear4* (*fea4*), a bZIP transcription factor with fasciated ears and tassels as well as greatly enlarged vegetative and inflorescence meristems (Pautler *et al*. 2015). QTL *TBN6.2* mapped to *tasselsheath1* (*tsh1*), a GATA class transcription factor that promotes bract growth and reduces branching (Whipple *et al*. 2010). QTL *TBN7.1* mapped to *ramosa1* (*ra1*), a C2H2 zinc-finger transcription factor which has tassels with an increased number of long branches as well as branched ears (Vollbrecht *et al*. 2005). QTL *TBN7.2* mapped to *tasselsheath4* (*tsh4*), a SBP-box transcription factor that functions to repress lateral organ growth and also affects phyllotaxy, axillary meristem initiation and meristem determinacy within the floral phase (Chuck *et al*. 2010). QTL *TBN8.1* mapped to *barren inflorescence1* (*bif1*), which shows a decreased production of branches and spikelet pairs (Barazesh and McSteen, 2008). QTL *TBN10.1* mapped to *zea floricaula leafy1* (*zfl1*), which together with its homolog *zfl2*, leads to a disruption of floral organ identity and patterning, as well as to defects in inflorescence architecture and in the vegetative to reproductive phase transition (Bomblies *et al*. 2003).

#### QTL for domestication traits

TILN is a classical domestication trait that measures difference in plant architecture between maize and its wild relative, teosinte – that is the low apical dominance of a highly branched teosinte plant as compared to the less-branched maize plant. We detected 18 small-effect QTLs that explained 68% of the total variance for TILN (Figure S32). Among them, QTL *TILN1.3* mapped to *tb1*, a TCP family of transcriptional regulators contributing to the increase in apical dominance during maize domestication (Doebley *et al*. 1997). Additionally, QTL *TILN3.2* mapped to *Zea AGAMOUS homolog2* (*zag2*), a MADS box gene recently found to be downstream of *tb1* (Studer *et al*. 2017). QTL *TILN1.1* and *TILN5.2* mapped to *zmm20* and *zmm26*, two other MADS box genes that were possible targets of selection during domestication (Zhao *et al*. 2011). QTL *TILN7.1* mapped to *PROSTRATE GROWTH1* (*PROG1*), a C2H2 zinc finger protein controlling a key change during rice domestication from prostrate to erect growth, and also affects plant architecture and yield-related traits (Jin *et al*., 2008; Tan *et al*. 2008). There are 13 genes in the support interval and the QTL peak is closest to *PROG1*, being ~14 kb 5’ of the start site (Figure S32). This is the first evidence that *PROG1* may have had a role in maize domestication.

GLUM is classical maize domestication trait measuring the dramatic change from the fruitcase-enveloped kernels of the teosinte ear to naked grains of maize ear. Previously, this trait was shown to be largely controlled by a single gene which is known as *teosinte glume architecture1* (*tga1*) (Wang *et al*. 2005). Interestingly, *tga1* is a direct target of *tb1*. We detected 11 QTLs that explained 62% of the total variance for GLUM. These QTL include a large effect QTL at *tga1* itself plus many small effect QTLs (Figure S33). Among the small effect QTL, our results show that two of them (*GLUM2.2* and *GLUM7.1*) mapped to MADS genes. In this regard, Studer *et al*. (2017) recently defined a maize domestication gene network in which *tga1* regulates multiple MADS-box transcription factors.

PROL is also an important domestication trait that measures difference in prolificacy or the many-eared plants of teosinte and the few-eared (one or two) plants of maize. Previously, a large effect QTL was fine-mapped to a region 2.7 kb upstream of *grassy tillers1* (*gt1*) (Wills *et al*. 2013). Interestingly, *gt1* is a known target of *tb1* (Whipple *et al*. 2011). We detected four QTLs that explained 39% of the total variance for PROL, which include a single large effect QTL plus three small effect QTLs (Figure S34). Concordantly, QTL *PROL1.1* mapped to *gt1*. QTL *PROL2.1* mapped to *zea floricaula leafy2* (*zfl2*), which was shown to have a pleiotropic effect on lateral branch number (Bomblies and Doebley, 2006). QTL *PROL3.1* mapped to *sparse inflorescence1* (*spi1*), a mutant that has defects in the initiation of axillary meristems and lateral organs during vegetative and inflorescence development in maize (Gallavotti *et al*. 2008). QTL *PROL5.1* mapped to *yabby9* (*yab9*), a class of transcription factor that might play important roles during maize domestication.

STAM measures the proportion of the terminal lateral inflorescence on the uppermost lateral branch that is staminate. Relative to domestication, this trait represents the sexual conversion of the terminal lateral inflorescence from tassel (staminate) in teosinte to ear (pistillate) in maize. Currently, *teosinte branched1* (*tb1*) and *tassel replace upper ears1* (*tru1*) are the only two genes that have been shown to regulate this sexual difference. We detected five QTLs that explained 27% of the total variance for STAM (Figure 4). QTL *STAM1.2* mapped to *tb1*, which is an important domestication gene known for various traits (Doebley *et al*. 1995). QTL *STAM3.1* mapped to *tru1* which is a direct target of *tb1* (Dong *et al*. 2017). QTL *STAM1.1* mapped to *tassel seed2* (*ts2*), a recessive mutant that produces pistillate spikelets in the terminal inflorescence (tassel) (Irish and Nelson, 1993). QTL *STAM3.2* mapped to *Zea mays MADS16* (*zmm16*), which shows high expression in tassel and silk. QTL *STAM7.1* mapped to *tassel sheath4* (*tsh4*), a SQUAMOSA PROMOTER BINDING (SBP)-box transcription factor that regulates the differentiation of lateral primordia (Chuck *et al*. 2010). In addition to these QTLs, two other STAM QTLs were detected by GWAS. Notably, a QTL on chromosome 1 (AGPv4 chr1:234.4-249.9Mb) is located upstream of *tb1* and co-localized with *STAM1.1* from a recent study (Yang *et al*. 2018). The known gene *anther ear1* (*an1*) is a strong candidate gene for this QTL since loss of *an1* function results in the development of staminate flowers in the ears (Bensen *et al*. 1995). The *tb1* QTL region was also detected by GWAS with a strong signal for interval AGPv4 chr1:264.1-283.1Mb.

**Figure 4.**
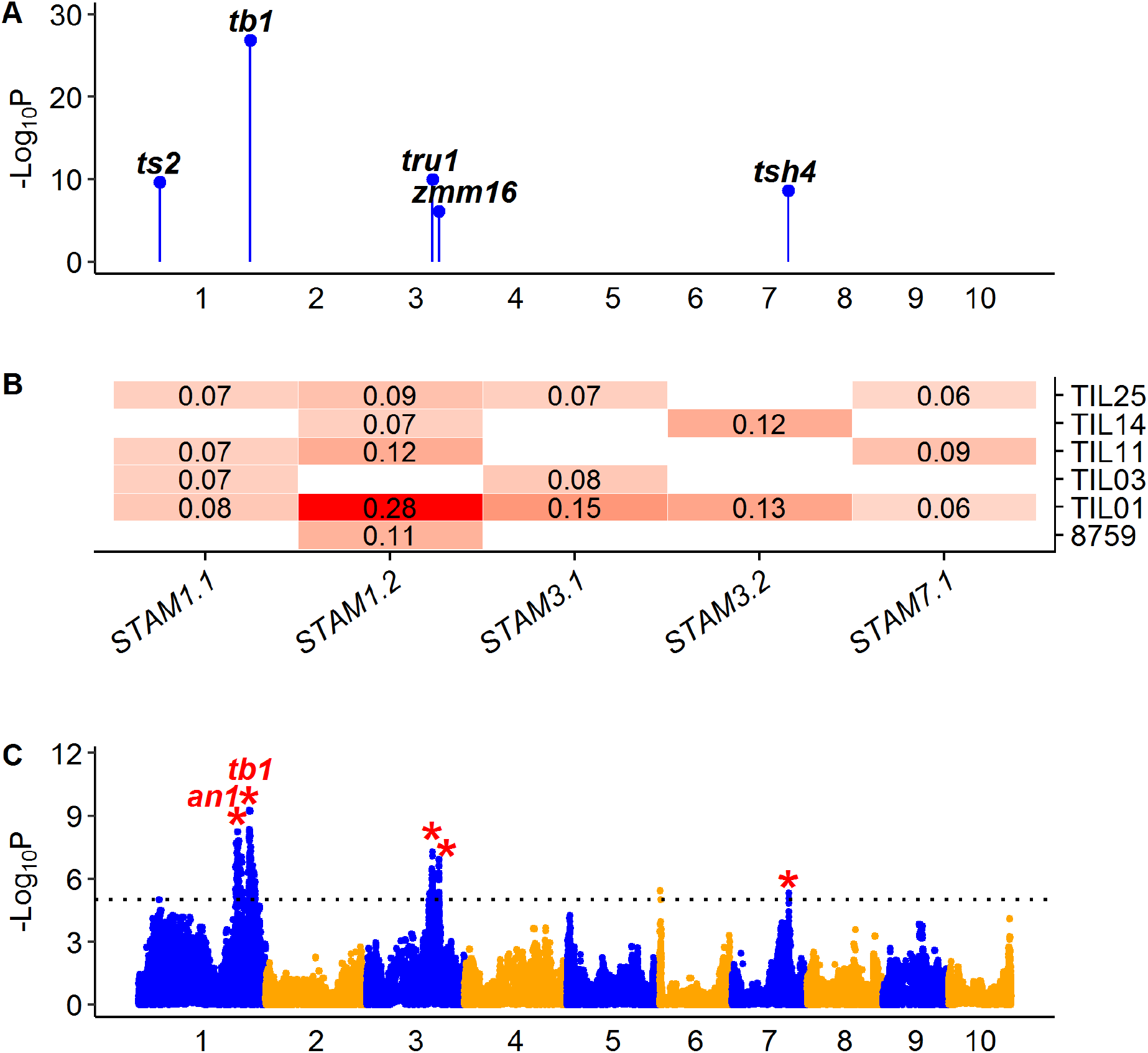
QTL characterization for domestication trait STAM. (A) Genomic distribution of five QTLs for STAM detected by JLM. The known candidate genes are shown above the corresponding QTLs in bold italic. (B) Heat map shows additive allele effects of teosinte relative to maize for five QTLs detected by JLM. The allele effect of teosinte parent 8759 was estimated from the 866 maize-teosinte BC_2_S_3_ RILs (Shannon 2012). Insignificant effects are shown as blank. The teosinte genotypes at all QTLs consistently contribute to a staminate lateral inflorescence. (C) Manhattan plot shows QTLs detected by GWAS. The significance threshold at LOD=5 is indicated by black dotted line. The red stars indicate GWAS signals overlapping with QTLs by JLM. In (A) and (C), chromosomes in odd and even numbers are shown in blue and orange colors, respectively.

SHN measures ear shattering, the loss of which is a key step during crop domestication (Doebley, 2006). Teosinte ears have abscission layers between the fruitcases (modified internodes) that allow the ear to shatter into single-seed units (fruitcase) at maturity. The maize ear lacks abscission layers and remains intact at maturity. Currently, only two maize orthologs (*ZmSh1-1* and *ZmSh1-5.1*+*ZmSh1-5.2*) of sorghum and rice *Shattering1* (*Sh1*) were verified for seed shattering (Lin *et al*. 2012). We detected six QTLs that explained 30% of the total variance for SHN (Figure S35). QTL *SHN1.1* and *SHN5.1* mapped to *Sh1.1* and *Sh1-5.1/5.2*, respectively, confirming prior identification of these maize paralogs of the sorghum shattering gene as strong candidates for our QTL.

KRN is a domestication trait measuring the dramatic change from the two-ranked teosinte ear to multiple (4 or more) ranked maize ear. We detected 24 small-effect QTLs that explained 62% of the total variance for KRN (Figure S36). Among them, QTL *KRN1.3* mapped to *indeterminate spikelet1* (*ids1*), an APETALA2-like transcription factor that specifies determinate fates by suppressing indeterminate growth within the spikelet meristem (Chuck *et al*. 1998). A previous fine-mapping study of KRN using a maize-teosinte BC_2_S_3_ RIL population also identified *ids1* is a strong candidate for KRN (Calderón *et al*. 2016). QTL *KRN4.2* mapped to *unbranched3* (*ub3*), a SBP transcription factor that has been shown to regulate kernel row number in both mutant and QTL studies (Chuck *et al*. 2014; Liu *et al*. 2015).

REPE for reddish-brownish pericarp is a trait that distinguishes teosinte kernels from those of most maize. The role of pigmentation in domestication is complex in that pigment can provide defense against molding and bird predation but can also impart bitterness and astringency (Morohashi *et al*. 2012). The red (or reddish brown) pigmentation often results from the accumulation of phlobaphenes - a flavonoid pigment (Morohashi *et al*. 2012). In the absence of the reddish-brown pigment, the kernels are white kernels unless anthocyanins (blue-purple) or carotenoids (yellow-orange) are present. Our results show that QTL *REPE1.1* mapped to *Pericarp color1* (*P1*) (Figure S37), which encodes an R2R3 Myb-like transcription factor that governs the biosynthesis of brick-red flavonoid pigments (Grotewold *et al*. 1994).

Results for 13 additional traits (ASI, BARE, CULM, DTS, EB, ED, EL, GLCO, KW, LFLN, LFWD, PLHT, and YEPE) are reported in supplemental figures and tables (Figure S30, S38-S49; Table S1).

### QTL detection and effects

To evaluate the power of QTL mapping using TeoNAM, we summarized the distribution of QTLs detected with significant effects in the different subpopulations. Among 255 QTLs for 22 traits, 246 QTLs (96%) were detected in two or more subpopulations, 186 QTLs (73%) were detected in three or more subpopulations, 83 QTLs (33%) were detected in four or more subpopulations and 29 QTLs (11%) were detected in all five subpopulations (Figure 5A). These percentages are conservative as not all traits were scored in all five subpopulations. If one considers whether the QTL was detected in subpopulations in which it was scored, then 205 QTLs (80%) were detected in at least half of the subpopulations and 39 QTLs (15%) were detected in all subpopulations.

**Figure 5.**
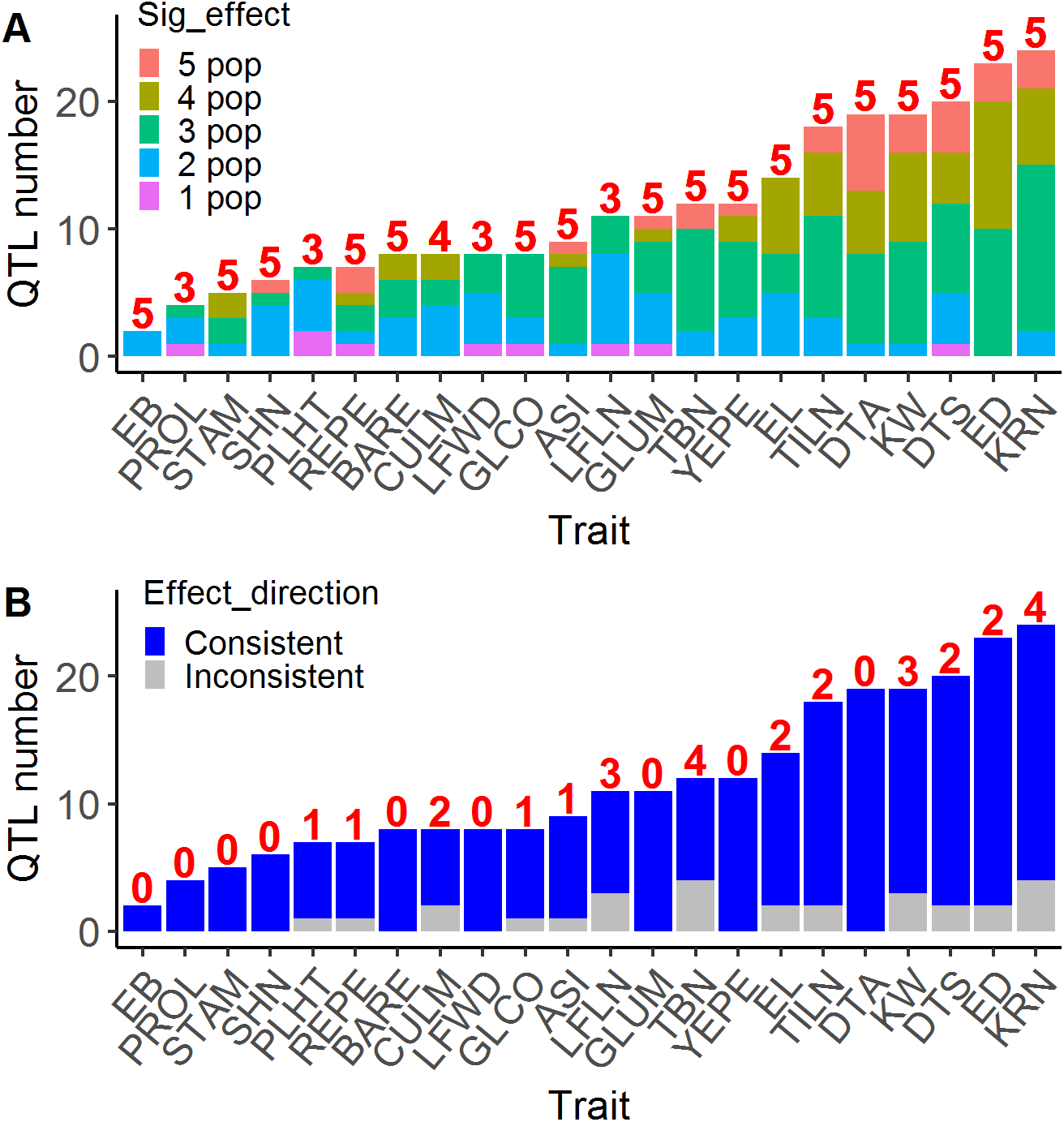
QTL detection and effects for all 22 traits. (A) Summary of QTL detection for all 22 traits. The number above the bar indicates the number of subpopulations in which the trait was scored. (B) Summary of QTL effect direction for all 22 traits. The number above the bar indicates the number of QTLs in which a teosinte allele was associated with the maize phenotype was detected.

The allelic effect from different teosinte parents were estimated simultaneously by JLM. For most QTLs, the allelic effects from different subpopulations are in the same direction (Figure 5B). For seven traits (EB, GLUM, LFWD, PROL, SHN, STAM, and YEPE), the teosinte genotypes were consistently associated with a teosinte phenotype and the W22 allele with a maize phenotype at all QTLs. For all other traits, cases in which a teosinte allele was associated with the maize phenotype were detected. For example, the teosinte genotype is associated with late flowering at most QTLs for DTA except *DTA5.2* and *DTA7.1*, for which the teosinte genotype consistently contributes to early flowering in at least three subpopulations (Figure 2). Similar results were observed for KRN and EL. The teosinte genotype is associated with fewer kernel row number (KRN) at most QTLs, but there is one QTL (*KRN5.1*) for which the teosinte genotype is consistently associated with more kernel row number in four subpopulations and also in the BC_2_S_3_ population (Figure S36). The teosinte genotype is associated with shorter ear length (EL) at most QTLs, but there are two QTLs (*EL4.1* and *EL9.1*) for which the teosinte genotype is consistently associated with longer ear length in four and two subpopulations, respectively (Figure S43). These QTLs might be worth exploring further for use in maize improvement.

We also observed some interesting results for different teosinte parents. For KW, the teosinte genotype from different subpopulations is associated with reduced kernel weight at most QTLs. Only three QTLs (*KW5.3*, *KW6.2* and *KW9.1*) are exceptions with one teosinte allele conferring heavier kernels. Interestingly for these three QTLs, the teosinte alleles with effects in the opposite direction are all from the TIL14 subpopulation (Figure S45). Similar results were observed for ED, where the teosinte genotype is associated with a decrease in ear diameter at most QTLs, but the teosinte allele from TIL03 at two QTLs (*ED2.1* and *ED6.1*) is associated with the increase of ear diameter (Figure S42). These results suggest that there are beneficial alleles from teosinte that could be utilized for maize improvement.

### Comparing and combining TeoNAM with the BC_2_S_3_

We compared TeoNAM with the previous maize-teosinte BC_2_S_3_ RIL population. The composite genetic map for TeoNAM is 1540 cM in length. The individual genetic maps based on the five subpopulations have an average length of 1461 cM with a range of 1348-1506 cM. The genetic map for BC_2_S_3_ RIL population is 1478 cM in length. Thus, the TeoNAM subpopulations are similar to the BC_2_S_3_ RIL population in genetic map length. The median length of homozygous teosinte segment in TeoNAM is 6 Mb. The median length of homozygous teosinte segment in BC_2_S_3_ population is 4.8 Mb. The longer segment length for TeoNAM is expected given it had one fewer generations of backcrossing and less opportunity for recombination. The mean number of homozygous teosinte segment in TeoNAM is 3502, and the number of homozygous teosinte segment in BC_2_S_3_ is 5745. The total length of teosinte segments for the five subpopulations is 67 GB (W22×TIL01), 87 GB (W22×TIL03), 66 GB (W22×TIL11), 56 GB (W22×TIL14) and 79 GB (W22×TIL25), and the BC_2_S_3_ (W22×8759) exceeds this range with 110 GB.

Previously, Shannon (2012) performed a comprehensive interval QTL analysis for 16 agronomic traits in the BC_2_S_3_ population and identified 218 QTLs for 16 traits. Among these traits, 14 traits were also scored in TeoNAM population. For the common 14 traits, 168 and 179 QTLs were detected for TeoNAM and BC_2_S_3_ population, respectively. The mean QTL support interval across 14 traits for BC_2_S_3_ is 5.7Mb, which is significantly smaller than TeoNAM of 17.2Mb (P=2.6E-08) (Figure S50). Among these QTLs, 50 QTLs overlapped between the two populations. For the common QTLs, the mean variance explained by QTL is 3.4% and 2.9% for BC_2_S_3_ and TeoNAM, respectively. Thus, there is no significant difference in QTL effect size (P=0.3) (Figure S51).

To maximize the power to detect QTLs, we combined TeoNAM and BC_2_S_3_ for eight traits (DTA, ED, EL, KRN, KW, GLCO, GLUM, and TILN) that were measured in all six subpopulations by the exactly same method to perform JLM. Before analysis, we imputed the genotype for BC_2_S_3_ at 4578 TeoNAM SNPs according to the flanking markers using the same procedure as for TeoNAM and permuted a new p-value cutoff for statistical significance for each trait. The LSMs from previous analysis (Shannon 2012) were used for JLM. With the combined TeoNAM-BC_2_S_3_ data, we detected 184 QTLs for these eight traits, which include 109 QTLs overlapped with TeoNAM, 80 QTLs overlapped with the BC_2_S_3_ and 32 novel QTLs not detected in either TeoNAM or the BC_2_S_3_ (Table S4). The QTLs with significant allele effects in multiple subpopulations will be good targets for fine-mapping. For future analysis of additional traits, one could combine TeoNAM and the BC_2_S_3_ together. The value of this combination is that there is one additional teosinte allele and increased QTL detection power, but the downside is that one would need to assay the BC_2_S_3_ population with 866 RILs plus TeoNAM with 1257 RILs.

## Discussion

RILs are powerful tools for dissecting complex genetic architecture of different traits and for gene discovery. RILs such as maize NAM population have been successfully used for genetic dissection of many traits (Buckler *et al*. 2009; Tian *et al*. 2011; Kump *et al*. 2011). RILs with the multiple parents greatly increase the power and precision to identify QTLs compared to the traditional bi-parent RIL population. Multi-parent RILs also enable the estimation of allele effects simultaneously from each inbred parent. Our TeoNAM RILs were created by crossing five teosinte inbred parents with a maize inbred parent, but differs from MaizeNAM in that we applied a generation of backcrossing to the maize parent before four generations of selfing. The power and precision of TeoNAM can be shown with several traits. For example, we detected 19 QTLs for DTA, among which many QTLs mapped to recently cloned genes such as *ZmCCT10*, *ZmCCT9*, *ZCN8*, *zagl1* and *ZmMADS69*. QTLs also mapped to some novel candidates such as *dlf1*, *si1*, *zag1*, *ZCN12*, *zmm19* and *zmm31*, which may have an important role in flowering time regulation.

For RIL populations, both JLM and GWAS are common methods for QTL detection. In this study, we identified 255 QTLs for 22 traits by JLM, and significant peaks were detected at 57 QTLs by GWAS, which suggests that GWAS is less powerful than JLM for mapping QTLs in TeoNAM. Nevertheless, there are a few instances in which GWAS gave evidence of closely linked QTL that were not separated by JLM. For example, we did not identify *an1*, a strong candidate for STAM QTL on chromosome 1 with JLM possibly because it’s closely linked to *tb1* (candidate of QTL *STAM1.1*), but we detected significant peaks at both *an1* and *tb1* through GWAS as it tests each SNP independently.

TeoNAM has allowed us to infer distinct genetic architectures for different traits. Traits like PROL and GLUM are controlled by a major effect QTL plus several QTLs of very small effect, while traits like DTA and KRN show more classic polygenic inheritance. These contrasting genetic architectures suggest that evolution during domestication did not always follow the same path. A variant of large effect at one locus with a few other small effect genes allowed naked kernels to evolve from covered kernels, but the more quantitative increase in the number of kernel rows required a larger number of genes with no single gene of substantially greater effect than all others.

In our study, a total of 15 domestication traits and 7 agronomic traits were analyzed. Further fine-mapping and gene cloning will be required to find the causal genes underlying QTLs for these traits. TeoNAM should also be useful for investigating genetic control of many new traits that we did not assay. Morphological traits such as root architecture, shoot apical meristem size, vasculature, and kernel shape can be explored. Also, molecular traits such as gene expression (eQTL), alternative splicing, grain protein content, and metabolites can also be explored to better understand the full spectrum of changes that occurred during maize domestication.

## Materials and Methods

### Population development

The teosinte NAM population was designed as a genetic resource for studying maize genetics and domestication. Five wild teosinte parents were chosen with four teosinte inbred lines that capture some diversity of *Zea mays* ssp. *parviglumis* (TIL01, TIL03, TIL11 and TIL14) and one teosinte inbred line of *Zea mays* ssp. *mexicana* (TIL25). The common parent is a modern maize inbred line W22 that has been widely used in maize genetics. The five teosinte parents were crossed to W22, and followed by one generation of backcross and four generations of selfing (Figure S1). We obtained 1257 BC_1_S_4_ recombinant inbred lines (RILs) with 223, 270, 219, 235 and 310 lines for W22×TIL01, W22×TIL03, W22×TIL11, W22×TIL14 and W22×TIL25, respectively.

### Marker Data

All DNA samples of 1257 lines were genotyped using Genotype-by-Sequencing (GBS) technology (Elshire *et al*. 2011). The genotypes were called from GBS raw sequencing reads using the TASSEL5-GBS Production Pipeline based on 955,690 SNPs in the ZeaGBSv2.7 Production TagsOnPhysicalMap file (Glaubitz *et al*. 2014). Then, the raw GBS markers were filtered in each RIL subpopulation using following steps. We first removed sites with minor allele frequencies below 5% and thinned sites with 64 bp apart using “Thin Sites by Position” in TASSEL5 (Bradbury *et al*. 2007), and then we ran FSFHap Imputation in TASSEL5 separately for each chromosome using the following parameters: backcross (bc), Phet=0.03125, Fillgaps=TRUE, and the default settings for other features. The imputed parental call files from the 10 chromosomes were then combined together and passed to R/qtl (Broman *et al*. 2003) to estimate genetic map. The B73 reference genome v2 was used to determine marker order, and genetic distances between markers was calculated using the Haldane mapping function as part of the *est.map* command with an assumed genotyping error rate of 0.001 taking the BC_1_S_4_ pedigree of the RIL into consideration (Shannon 2012). Bad genetic markers were identified by visual inspection of the genetic map and removed, then we repeated all filtering steps. Finally, an average of 13,733 high-quality SNPs was obtained for each subpopulation (Table 1).

### Field design and phenotyping

The teosinte NAM population was planted using a randomized complete block design at the University of Wisconsin West Madison Agricultural Research Station (UW - WMARS) in different years. The subpopulations W22×TIL01, W22×TIL03, W22×TIL11 were grown in summer 2015 and 2016, the subpopulation W22×TIL14 was grown in summer 2016 and 2017, and the subpopulation W22×TIL25 was grown in summer 2017 with two blocks. We planted one subpopulation within each block, and all lines were randomized within each block. Each row had 16 seeds planted with 1-foot apart, and spacing between any two rows was 30-inch.

Twenty-two traits were scored (Table 2): days to anthesis (DTA) (number of days between planting and when at least half the plants in a plot were shedding pollen); days to silk (DTA) (number of days between planting and when at least half the plants in a plot were showing silk); anthesis-silk interval (ASI) (number of days between anthesis and silk); tassel branch number (TBN) (number of tassel branches on the main stalk); culm diameter (CULM) (diameter of the narrowest plane of main stalk right above the ground); plant height (PLHT) (distance from ground to the topmost node on the main stalk); leaf length (LFLN) (length of a well-developed leaf, usually 4th-6th from top); leaf width (LFWD) (width of a well-developed leaf, usually 4th-6th from top); tiller number (TILN) (number of tillers surrounding main stalk); prolificacy (PROL) (0 vs. 1 for absence/presence of secondary ears at the topmost branch-bearing node on the main stalk); ear branch number (EB) (number of branch on the primary lateral inflorescence); staminate spikelet (STAM) (0-3 scale for spikelet sex on the primary lateral inflorescence, where 0 indicates completely feminized, and 3 indicates completely staminate); kernel row number (KRN) (number of internode columns on the primary lateral inflorescence); ear length (EL) (length of the primary lateral inflorescence); ear diameter (ED) (diameter of the primary lateral inflorescence); kernel weight (KW) (average weight of 50 random kernels from 5 ears); shattering (SHN) (number of pieces into which an ear shattered when dropped to the floor from a height of ~1.8m); barren ear base (BARE) (0-2 scale for lack of kernels at the base of ear, where 0 indicates kernels present at the base, and 2 indicates no developed kernels at the base of the ear); glume score (GLUM) (0-3 scale for glume size, where 0 indicates small and 3 indicates large); glume color (GLCO) (0-4 scale glume color for white through brown); red pericarp (REPE) (0-2 scale for colorless to red pericarp); yellow pericarp (YEPE) (0-2 scale for dull yellow to bright yellow pericarp). The average trait value from two years were used for QTL analysis.

### Genetic map construction and marker imputation

A composite genetic map was constructed for the TeoNAM population. The markers from the five RIL subpopulations were combined together into 51,544 unique SNPs, and the missing genotypes were imputed according to the flanking markers. If the flanking markers have same genotypes, the missing genotype was imputed as the same with flanking markers, otherwise left as missing. The imputed genotypes were then passed to R/qtl software to estimate the genetic map.

Since stepwise regression cannot use individuals with missing marker data, we performed a further step to impute missing data around break point as previously described (Tian *et al*. 2011). First, we transformed genotype to numeric format, in which markers with homozygous W22 parent were coded as 0, markers with homozygous non-W22 parent were coded as 2, and markers with heterozygous genotypes were coded as 1. Markers within breakpoint were imputed according to the genetic distance of flanking two markers. Considering stepwise regression is computationally intensive, we thinned SNPs within 0.1 cM. We finally obtained 4,578 markers for subsequent joint linkage analysis.

### Simple QTL mapping

QTL mapping was carried out using a modified version of R/qtl (Broman *et al*., 2003) which takes into account the BC_1_S_4_ pedigree of the RILs (Shannon, 2012). For each trait, a total of 1000 permutation tests were used to determine the significance threshold level for claiming QTLs. After permutation, an approximate LOD score of 4.0 at P < 0.05 was obtained across all traits. With the LOD threshold, simple interval mapping was first fitted using Haley-Knott regression implemented in the *scanone* command of R/qtl. The multiple QTL model was then applied to search for additional QTL and accurately refine QTL positions using *refineqtl* and *addqtl* in R/qtl. The entire process was repeated until significant QTLs could no longer be added. The total phenotypic variation explained by all QTLs was calculated from a full model that fitted all QTL terms in the model using the *fitqtl* function. The percentage of phenotypic variation explained by each QTL was estimated using a drop-one-ANOVA analysis implemented with the *fitqtl* function. The confidence interval for each QTL was defined using a 1.5-LOD support interval. To make results comparable among five subpopulations, the composite genetic map was used for QTL mapping.

### Joint linkage mapping

To map QTL in the TeoNAM population, a joint linkage mapping (JLM) procedure was performed as previously described (Buckler *et al*. 2009; Tian *et al*. 2011). First, a total of 1000 permutation were performed to determine the significance cutoff for each trait. JLM was performed using the stepwise linear regression fixed model implemented by PROC GLMSELECT procedure in SAS software. The family main effect was fit first, and then marker effects nested within families were selected to enter or leave the model based on the permutated P-value using a marginal F-test. After the model was fit with stepwise regression, each marker was dropped from the full model one at a time and a single best marker was refit to improve the overall fit of the model using the remaining QTL as background. A threshold of α=0.05 was used to declare significant allele effects across families within each QTL identified by stepwise regression. The QTL support interval was calculated by adding each marker from the same chromosome of that QTL at a time to the full model. If the p-value of the marginal F-test of the QTL was not significant at the 0.01 level, the flanking marker should be in the support interval for the QTL as the new flanking marker explained the QTL as well as the original marker.

### GWAS

A genome-wide association study (GWAS) approach was also used to map QTL in the TeoNAM population. Since GBS produces relatively low-density markers, the 955,690 raw SNPs from GBS pipeline were filtered using a less conservative criteria: MAF>0.01, missing rate < 0.75, and heterozygosity rate < 0.1. After this filtering, 181,404 GBS SNPs were used to run FSFHap Imputation in TASSEL5 separately for each chromosome and subpopulation using the following parameters: backcross (bc), Phet=0.03125, Fillgaps=TRUE, and the default settings for other features. Imputed genotypes were then combined together and SNPs with missing rate more than 0.2 and MAF less than 0.05 across 1257 RILs were removed and a total of 118,838 SNPs were kept and used for GWAS. GWAS was performed using a linear mixed model accounting for population structure (Q) and kinship matrix (K), where Q was computed as the first five principle components and K was calculated using centered IBS method as implemented in TASSEL (Bradbury *et al*. 2007). The P value below P=0.00001 (LOD=5) was considered as significance threshold following a previous study (Kremling *et al*. 2018).

### QTL candidate analysis

To report the QTL position following the latest genomic version, we used the CrossMap (Zhao *et al*. 2014) software to uplift the GBS SNP positions from maize B73 reference AGPv2 coordinates to AGPv4 coordinates. QTL candidates were analyzed by checking the gene annotations of genes within QTL support intervals.

### Data Availability

Seeds for all 1257 RILs are available through the Maize Genetics Cooperative Stock Center and the SNP genotypes of TeoNAM are available at the Cyverse Discovery Environment under the directory: /iplant/home/shared/panzea/genotypes/GBS/TeosinteNAM/. The genotypes were uploaded with AGPv2 position in the marker name.

## Acknowledgements

This research was supported by the US National Science Foundation (NSF) grants IOS 1238014 and China Postdoctoral Science Foundation (2018M640204). No conflict of interest declared. We thank Karl Broman for suggestions on the analyses, and Jesse Rucker, Elizabeth Buschert, Eric Rentmeester, Adam Mittermaier, David Sierakowski, and Brian Schaeffer for assistance with field work and phenotyping.

## Literature Cited

Ambrose, B. A., Lerner, D. R., Ciceri, P., Padilla, C. M., Yanofsky, M. F., and Schmidt, R. J. (2000). Molecular and genetic analyses of the *silky1* gene reveal conservation in floral organ specification between eudicots and monocots. Mol. Cell 5:569–579.

Bandillo, N., Raghavan, C., Muyco, P. A., Sevilla, M. A. L., Lobina, I. T., Dilla-Ermita, C. J., Tung, C. W., McCouch, S., Thomson, M., Mauleon, R., et al. (2013). Multi-parent advanced generation inter-cross (MAGIC) populations in rice: Progress and potential for genetics research and breeding. Rice 6:1–15.

Barazesh, S., and McSteen, P. (2008). *Barren inflorescence1* functions in organogenesis during vegetative and inflorescence development in maize. Genetics 179:389–401.

Bensen, R.J., Johal, G.S., Crane, V.C., Tossberg, J.T., Schnable, P.S., Meeley, R.B. and Briggs, S.P. (1995). Cloning and characterization of the maize *An1* gene. Plant Cell 7:75–84.

Bomblies, K., Wang, R.L., Ambrose, B.A., Schmidt, R.J., Meeley, R.B. and Doebley, J. (2003). Duplicate *FLORICAULA/LEAFY* homologs *zfl1* and *zfl2* control inflorescence architecture and flower patterning in maize. Development 130:2385–2395.

Bomblies, K., and Doebley, J. F. (2006). Pleiotropic effects of the duplicate maize *FLORICAULA/LEAFY* genes *zfl1* and *zfl2* on traits under selection during maize domestication. Genetics 172:519–531.

Bouchet, S., Olatoye, M. O., Marla, S. R., Perumal, R., and Tesso, T. (2017). Increased power to dissect adaptive traits in global sorghum diversity using a nested association mapping population. Genetics 206:573–585.

Bradbury, P. J., Zhang, Z., Kroon, D. E., Casstevens, T. M., Ramdoss, Y., and Buckler, E. S. (2007). TASSEL: software for association mapping of complex traits in diverse samples. Bioinformatics 23:2633–2635.

Broman, K. W., Wu, H., Sen, Ś., and Churchill, G. A. (2003). R/qtl: QTL mapping in experimental crosses. Bioinformatics 19:889–890.

Buckler, E.S., Holland, J.B., Bradbury, P.J., Acharya, C.B., Brown, P.J., Browne, C., Ersoz, E., Flint-Garcia, S., Garcia, A., Glaubitz, J.C., et al. (2009). The genetic architecture of maize flowering time. Science 325:714–718.

Calderón, C. I., Yandell, B. S., and Doebley, J. F. (2016). Fine mapping of a QTL associated with kernel row number on chromosome 1 of maize. PLoS One 11:e0150276.

Chuck, G., Meeley, R. B., and Hake, S. (1998). The control of maize spikelet meristem fate by the *APETALA2*-like gene *indeterminate spikelet1*. Genes Dev. 12:1145–1154.

Chuck, G., Whipple, C., Jackson, D., and Hake, S. (2010). The maize SBP-box transcription factor encoded by *tasselsheath4* regulates bract development and the establishment of meristem boundaries. Development 137:1243–1250.

Chuck, G. S., Brown, P. J., Meeley, R., and Hake, S. (2014). Maize *SBP-box* transcription factors *unbranched2* and *unbranched3* affect yield traits by regulating the rate of lateral primordia initiation. Proc. Natl. Acad. Sci. USA 111:18775–18780.

Churchill, G.A., Airey, D.C., Allayee, H., Angel, J.M., Attie, A.D., Beatty, J., Beavis, W.D., Belknap, J.K., Bennett, B., Berrettini, W., et al. (2004). The Collaborative cross, a communnity resource for the gentic analysis fo complex traits. Nat. Genet. 36:1133–1137.

Dell’Acqua, M., Gatti, D. M., Pea, G., Cattonaro, F., Coppens, F., Magris, G., Hlaing, A. L., Aung, H. H., Nelissen, H., Baute, J., et al. (2015). Genetic properties of the MAGIC maize population: a new platform for high definition QTL mapping in Zea mays. Genome Biol. 16:1–23.

Doebley, J. F., Gaut, B. S., and Smith, B. D. (2006). The molecular genetics of crop domestication. Cell 127:1309–1321.

Doebley, J., Stec, A. and Gustus, C. (1995). *teosinte branched1* and the origin of maize: evidence for epistasis and the evolution of dominance. Genetics 141:333–346.

Doebley, J., Stec, A. and Hubbard, L. (1997). The evolution of apical dominance in maize. Nature 386:485–488.

Dong, Z., Li, W., Unger-Wallace, E., Yang, J., Vollbrecht, E. and Chuck, G. (2017). Ideal crop plant architecture is mediated by *tassels replace upper ears1*, a BTB/POZ ankyrin repeat gene directly targeted by TEOSINTE BRANCHED1. Proc. Natl. Acad. Sci. USA 114:E8656–E8664.

Elshire, R. J., Glaubitz, J. C., Sun, Q., Poland, J. A., Kawamoto, K., Buckler, E. S., and Mitchell, S. E. (2011). A robust, simple genotyping-by-sequencing (GBS) approach for high diversity species. PLoS One 6:e19379.

Evans, M. M. S., and Kermicle, J. L. (2001). *Teosinte crossing barrier1*, a locus governing hybridization of teosinte with maize. Theor. Appl. Genet. 103:259–265.

Fragoso, C. A., Moreno, M., Wang, Z., Heffelfinger, C., Arbelaez, L. J., Aguirre, J. A., Franco, N., Romero, L. E., Labadie, K., Zhao, H., et al. (2017). Genetic architecture of a rice nested association mapping population. G3 (Bethesda) 7:1913–1926.

Gallavotti, A., Barazesh, S., Malcomber, S., Hall, D., Jackson, D., Schmidt, R. J., and McSteen, P. (2008). *sparse inflorescence1* encodes a monocot-specific *YUCCA*-like gene required for vegetative and reproductive development in maize. Proc. Natl. Acad. Sci. USA 105:15196–15201.

Glaubitz, J. C., Casstevens, T. M., Lu, F., Harriman, J., Elshire, R. J., Sun, Q., and Buckler, E. S. (2014). TASSEL-GBS: A high capacity genotyping by sequencing analysis pipeline. PLoS One 9:e90346.

Grotewold, E., Drummond, B.J., Bowen, B. and Peterson, T. (1994). The *myb*-homologous *P* gene controls phlobaphene pigmentation in maize floral organs by directly activating a flavonoid biosynthetic gene subset. Cell 76:543–553.

Guo, L., Wang, X., Zhao, M., Huang, C., Li, C., Li, D., Yang, C. J., York, A. M., Xue, W., Xu, G., et al. (2018). Stepwise *cis*-regulatory changes in *ZCN8* contribute to maize flowering-time adaptation. Curr. Biol. 28:3005–3015.

Huang, B. E., George, A. W., Forrest, K. L., Kilian, A., Hayden, M. J., Morell, M. K., and Cavanagh, C. R. (2012). A multiparent advanced generation inter-cross population for genetic analysis in wheat. Plant Biotechnol. J. 10:826–839.

Huang, C., Chen, Q., Xu, G., Xu, D., Tian, J. and Tian, F. (2016). Identification and fine mapping of quantitative trait loci for the number of vascular bundle in maize stem. J. Integr. Plant Biol. 58:81–90.

Huang, C., Sun, H., Xu, D., Chen, Q., Liang, Y., Wang, X., Xu, G., Tian, J., Wang, C., Li, D., et al. (2017). *ZmCCT9* enhances maize adaptation to higher latitudes. Proc. Natl. Acad. Sci. USA 115:E334–E341.

Hung, H.-Y., Shannon, L. M., Tian, F., Bradbury, P. J., Chen, C., Flint-Garcia, S. A., McMullen, M. D., Ware, D., Buckler, E. S., Doebley, J. F., et al. (2012). *ZmCCT* and the genetic basis of day-length adaptation underlying the postdomestication spread of maize. Proc. Natl. Acad. Sci. USA 109:E1913–E1921.

Irish, E.E. and Nelson, T.M. (1993). Development of *tassel seed 2* inflorescences in maize. Am. J. Bot. 80:292–299.

Jin, J., Huang, W., Gao, J.P., Yang, J., Shi, M., Zhu, M.Z., Luo, D. and Lin, H.X. (2008). Genetic control of rice plant architecture under domestication. Nat. Genet. 40:1365–1369.

Jordan, K. W., Wang, S., He, F., Chao, S., Lun, Y., Paux, E., Sourdille, P., Sherman, J., Akhunova, A., Blake, N. K., et al. (2018). The genetic architecture of genome-wide recombination rate variation in allopolyploid wheat revealed by nested association mapping. Plant J. 95:1039–1054.

King, E. G., Merkes, C. M., McNeil, C. L., Hoofer, S. R., Sen, S., Broman, K. W., Long, A. D., and Macdonald, S. J. (2012). Genetic dissection of a model complex trait using the *Drosophila* Synthetic Population Resource. Genome Res. 22:1558–1566.

Kover, P. X., Valdar, W., Trakalo, J., Scarcelli, N., Ehrenreich, I. M., Purugganan, M. D., Durrant, C., and Mott, R. (2009). A multiparent advanced generation inter-cross to fine-map quantitative traits in *Arabidopsis thaliana*. PLoS Genet. 5:e1000551.

Kremling, K.A., Chen, S.Y., Su, M.H., Lepak, N.K., Romay, M.C., Swarts, K.L., Lu, F., Lorant, A., Bradbury, P.J. and Buckler, E.S. (2018). Dysregulation of expression correlates with rare-allele burden and fitness loss in maize. Nature 555:520–523.

Kump, K. L., Bradbury, P. J., Wisser, R. J., Buckler, E. S., Belcher, A. R., Oropeza-Rosas, M. A., Zwonitzer, J. C., Kresovich, S., McMullen, M. D., Ware, D., et al. (2011). Genome-wide association study of quantitative resistance to southern leaf blight in the maize nested association mapping population. Nat. Genet. 43:163–168.

Lang, Z., Wills, D. M., Lemmon, Z. H., Shannon, L. M., Bukowski, R., Wu, Y., Messing, J., and Doebley, J. F. (2014). Defining the role of *prolamin-box binding factor1* gene during maize domestication. J. Hered. 105:576–582.

Leiboff, S., DeAllie, C.K. and Scanlon, M.J. (2016). Modeling the morphometric evolution of the maize shoot apical meristem. Front. Plant Sci. 7:1651.

Liang, Y., Liu, Q., Wang, X., Huang, C., Xu, G., Hey, S., Lin, H. Y., Li, C., Xu, D., Wu, L., et al. (2019). *ZmMADS69* functions as a flowering activator through the *ZmRap2.7*-*ZCN8* regulatory module and contributes to maize flowering time adaptation. New Phytol. 221:2335–2347.

Li, D., Wang, X., Zhang, X., Chen, Q., Xu, G., Xu, D., Wang, C., Liang, Y., Wu, L., Huang, C., et al. (2016). The genetic architecture of leaf number and its genetic relationship to flowering time in maize. New Phytol. 210:256–268.

Lin, Z., Li, X., Shannon, L. M., Yeh, C. T., Wang, M. L., Bai, G., Peng, Z., Li, J., Trick, H. N., Clemente, T. E., et al. (2012). Parallel domestication of the *Shattering1* genes in cereals. Nat. Genet. 44:720–724.

Liu, L., Du, Y., Shen, X., Li, M., Sun, W., Huang, J., Liu, Z., Tao, Y., Zheng, Y., Yan, J., et al. (2015). *KRN4* controls quantitative variation in maize kernel row number. PLoS Genet. 11:e1005670.

Mackay, I. J., Bansept-basler, P., Barber, T., Bentley, A. R., Cockram, J., Gosman, N., Horsnell, R., Howells, R., Sullivan, D. M. O., Rose, G. A., et al. (2014). An eight-parent multiparent advanced generation inter-cross population for winter-sown wheat: creation, properties, and validation. G3 (Bethesda) 4:1603–1610.

Maurer, A., Draba, V., Jiang, Y., Schnaithmann, F., Sharma, R., Schumann, E., Kilian, B., Reif, J. C., and Pillen, K. (2015). Modelling the genetic architecture of flowering time control in barley through nested association mapping. BMC Genomics 16:1–12.

Meng, X., Muszynski, M. G., and Danilevskaya, O. N. (2011). The *FT*-like *ZCN8* gene functions as a floral activator and is involved in photoperiod sensitivity in maize. Plant Cell 23:942–960.

Morohashi, K., Casas, M. I., Ferreyra, M. L. F., Mejía-Guerra, M. K., Pourcel, L., Yilmaz, A., Feller, A., Carvalho, B., Emiliani, J., Rodriguez, E., et al. (2012). A genome-wide regulatory framework identifies maize *Pericarp Color1* controlled genes. Plant Cell 24:2745–2764.

Nice, L. M., Steffenson, B. J., Brown-Guedira, G. L., Akhunov, E. D., Liu, C., Kono, T. J. Y., Morrell, P. L., Blake, T. K., Horsley, R. D., Smith, K. P., et al. (2016). Development and genetic characterization of an advanced backcross-nested association mapping (AB-NAM) population of wild × cultivated barley. Genetics 203:1453–1467.

Pautler, M., Eveland, A. L., LaRue, T., Yang, F., Weeks, R., Lunde, C., Je, B. Il, Meeley, R., Komatsu, M., Vollbrecht, E., et al. (2015). *FASCIATED EAR4* encodes a bZIP transcription factor that regulates shoot meristem size in maize. Plant Cell 27:104–120.

Schmidt, R.J., Veit, B., Mandel, M.A., Mena, M., Hake, S. and Yanofsky, M.F. (1993). Identification and molecular characterization of *ZAG1*, the maize homolog of the Arabidopsis floral homeotic gene *AGAMOUS*. Plant Cell 5:729–737.

Shannon, L.M. (2012). The genetic architecture of maize domestication and range expansion. [PhD Dissertation] The University of Wisconsin-Madison.

Studer, A. J., Wang, H., and Doebley, J. F. (2017). Selection during maize domestication targeted a gene network controlling plant and inflorescence architecture. Genetics 207:755–765.

Tan, L., Li, X., Liu, F., Sun, X., Li, C., Zhu, Z., Fu, Y., Cai, H., Wang, X., Xie, D., et al. (2008). Control of a key transition from prostrate to erect growth in rice domestication. Nat. Genet. 40:1360–1364.

Theissen, G., Becker, A., Di Rosa, A., Kanno, A., Kim, J.T., Münster, T., Winter, K.U. and Saedler, H. (2000). A short history of MADS-box genes in plants. Plant Molecular Evolution pp.115–149. Springer, Dordrecht.

Tian, F., Bradbury, P. J., Brown, P. J., Hung, H., Sun, Q., Flint-Garcia, S., Rocheford, T. R., McMullen, M. D., Holland, J. B., and Buckler, E. S. (2011). Genome-wide association study of leaf architecture in the maize nested association mapping population. Nat. Genet. 43:159–162.

Vollbrecht, E., Springer, P. S., Goh, L., Buckler IV, E. S., and Martienssen, R. (2005). Architecture of floral branch systems in maize and related grasses. Nature 436:1119–1126.

Wang, H., Nussbaum-Wagler, T., Li, B., Zhao, Q., Vigouroux, Y., Faller, M., Bomblies, K., Lukens, L., and Doebley, J. F. (2005). The origin of the naked grains of maize. Nature 436:714–719.

Wang, X., Chen, Q., Wu, Y., Lemmon, Z. H., Xu, G., Huang, C., Liang, Y., Xu, D., Li, D., Doebley, J. F., et al. (2018). Genome-wide analysis of transcriptional variability in a large maize-teosinte population. Mol. Plant 11:443–459.

Whipple, C. J., Hall, D. H., DeBlasio, S., Taguchi-Shiobara, F., Schmidt, R. J., and Jackson, D. P. (2010). A conserved mechanism of bract suppression in the grass family. Plant Cell 22:565–578.

Whipple, C.J., Kebrom, T.H., Weber, A.L., Yang, F., Hall, D., Meeley, R., Schmidt, R., Doebley, J., Brutnell, T.P. and Jackson, D.P. (2011). *grassy tillers1* promotes apical dominance in maize and responds to shade signals in the grasses. Proc. Natl. Acad. Sci. USA 108:E506–E512.

Wills, D. M., Fang, Z., York, A. M., Holland, J. B., and Doebley, J. F. (2017). Defining the role of the MADS-box gene, *Zea Agamous-like1*, a target of selection during maize domestication. J. Hered. 109:333–338.

Wills, D. M., Whipple, C. J., Takuno, S., Kursel, L. E., Shannon, L. M., Ross-Ibarra, J., and Doebley, J. F. (2013). From many, one: genetic control of prolificacy during maize domestication. PLoS Genet. 9:e1003604.

Xavier, A., Jarquin, D., Howard, R., Ramasubramanian, V., Specht, J. E., Graef, G. L., Beavis, W. D., Diers, B. W., Song, Q., Cregan, P., et al. (2018). Genome-wide analysis of grain yield stability and environmental interactions in a multiparental soybean population. G3 (Bethesda) 8:519–529.

Xiao, Y., Tong, H., Yang, X., Xu, S., Pan, Q., Qiao, F., Raihan, M. S., Luo, Y., Liu, H., Zhang, X., et al. (2016). Genome-wide dissection of the maize ear genetic architecture using multiple populations. New Phytol. 210:1095–1106.

Xu, D., Wang, X., Huang, C., Xu, G., Liang, Y., Chen, Q., Wang, C., Li, D., Tian, J., Wu, L., et al. (2017a). *Glossy15* plays an important role in the divergence of the vegetative transition between maize and its progenitor, teosinte. Mol. Plant 10:1579–1583.

Xu, G., Wang, X., Huang, C., Xu, D., Li, D., Tian, J., Chen, Q., Wang, C., Liang, Y., Wu, Y., et al. (2017b). Complex genetic architecture underlies maize tassel domestication. New Phytol. 214:852–864.

Yang, C.J. (2018). Dissection of the genetic architecture of domestication traits in maize and its ancestor teosinte. [PhD Dissertation] The University of Wisconsin-Madison.

Yang, Q., Li, Z., Li, W., Ku, L., Wang, C., Ye, J., Li, K., Yang, N., Li, Y., Zhong, T., et al. (2013). CACTA-like transposable element in *ZmCCT* attenuated photoperiod sensitivity and accelerated the postdomestication spread of maize. Proc. Natl. Acad. Sci. USA 110:16969–16974.

Yu, J., Holland, J. B., McMullen, M. D., and Buckler, E. S. (2008). Genetic design and statistical power of nested association mapping in maize. Genetics 178:539–551.

Zhang, Z., Zhang, X., Lin, Z., Wang, J., Xu, M., Lai, J., Yu, J., and Lin, Z. (2018). The genetic architecture of nodal root number in maize. Plant J. 93:1032–1044.

Zhao, Q., Weber, A. L., McMullen, M. D., Guill, K., and Doebley, J. (2011). MADS-box genes of maize: frequent targets of selection during domestication. Genet. Res. 93:65–75.

Zhao, H., Sun, Z., Wang, J., Huang, H., Kocher, J. P., and Wang, L. (2014). CrossMap: a versatile tool for coordinate conversion between genome assemblies. Bioinformatics 30:1006–1007.

